# Assortment of flowering time and immunity alleles in natural *Arabidopsis thaliana* populations suggests immunity and vegetative lifespan strategies coevolve

**DOI:** 10.1101/131136

**Authors:** Shirin Glander, Fei He, Gregor Schmitz, Anika Witten, Arndt Telschow, J. de Meaux

## Abstract

The selective impact of pathogen epidemics on host defenses can be strong but remains transient. By contrast, life-history shifts can durably and continuously modify the balance between costs and benefits of immunity, which arbitrates the evolution of host defenses. Their impact on the evolutionary dynamics of host immunity, however, has seldom been documented. Optimal investment into immunity is expected to decrease with shortening lifespan, because a shorter life decreases the probability to encounter pathogens or enemies. Here, we document that in natural populations of *Arabidopsis thaliana*, the expression levels of immunity genes correlate positively with flowering time, which in annual species is a proxy for lifespan. Using a novel genetic strategy based on bulk-segregants, we partitioned flowering time-dependent from – independent immunity genes and could demonstrate that this positive co-variation can be genetically separated. It is therefore not explained by the pleiotropic action of some major regulatory genes controlling both immunity and lifespan. Moreover, we find that immunity genes containing variants reported to impact fitness in natural field conditions are among the genes whose expression co-varies most strongly with flowering time. Taken together, these analyses reveal that natural selection has likely assorted alleles promoting lower expression of immunity genes with alleles that decrease the duration of vegetative lifespan in *A. thaliana* and *vice versa*. This is the first study documenting a pattern of variation consistent with the impact that selection on flowering time is predicted to have on diversity in host immunity.

## INTRODUCTION

The ability of organisms to defend against pathogens is a major determinant of survival in natural populations (Parker & Gilbert, 2004; Chisholm *et al.*, 2006; Lee & Mazmanian, 2010). Pathogens have long been suspected to impose a fast evolution of the host immune system and the “Red Queen” Hypothesis is nowadays a keystone of evolutionary biology (Van Valen, 1973; Liow *et al.*, 2011). Evidence that pathogens drive the molecular evolution of host defense systems has been accumulating in an array of plant and animal systems (Bergelson *et al.*, 2001; de Meaux & Mitchell-Olds, 2003; Moeller & Tiffin, 2005; Ravensdale *et al.*, 2010; Laine *et al.*, 2010; Maekawa *et al.*, 2011; Dybdahl *et al.*, 2014; Karasov *et al.*, 2014; Siddle & Quintana-Murci, 2014; Parker *et al.*, 2015; Metzger *et al.*, 2016).

Yet, the possible impact of changes in ecology on the evolution of defense systems should also be considered as they may durably change the exposure of hosts to pathogens. Invasive species, for example, owe much of their success to the release from pathogen and pest pressures (Mitchell & Power, 2003; Mitchell *et al.*, 2010). Similarly, shifts in life history can alter the balance between costs and benefits of host defense systems (Herms & Mattson, 1992). Shifting from perennial to annual life cycles, or evolving from a winter-annual to summer-annual cycling occurs frequently across plant phylogenies (Garnier, 1992; Michaels *et al.*, 2003; Franks *et al.*, 2007; Tank & Olmstead, 2008; Matthew Ogburn & Edwards, 2015; Kiefer *et al.*, 2017). The reduction in lifespan that follows such life history changes concomitantly reduces the probability to encounter ennemies (Jokela *et al.*, 2000). As a matter of fact, woody plant species with longer lifespan often display stronger herbivore defenses (Endara & Coley, 2010). As a consequence, immunity and lifespan are expected to coevolve.

*Arabidopsis thaliana* populations offer an optimal model for catching the co-evolution of life history and immunity in the act. *A. thaliana* has become over the last decade a powerful model system to address ecological questions at the genetic level (Mitchell-Olds & Schmitt, 2006; Bergelson & Roux, 2010; Roux & Bergelson, 2016). Experiments in common gardens have been performed to describe the architecture of natural variation in fitness and to infer geographic distributions of locally adaptive mutations (Fournier-Level *et al.*, 2011; Hancock *et al.*, 2011; Fournier-Level *et al.*, 2016). Analyses of mutants and recombinant inbred lines (RIL) have allowed reconstructing the contribution of phenotypes to fitness (Wilczek *et al.*, 2009; Chiang *et al.*, 2013; Fournier-Level *et al.*, 2013). Secondary chemical compounds were shown to have evolved to deter predominant herbivores in natural populations (Brachi *et al.*, 2013; Kerwin *et al.*, 2015). Clinal variation along the latitudinal range of the species reveals how phenotypes expressed along the life cycle are jointly shaped by natural selection (Lasky, 2012; Debieu *et al.*, 2013; Vidigal *et al.*, 2016).

*A. thaliana* is arguably one of the species for which we have the largest amount of genetic and phenotypic information on both immune reactions against pathogens and variation in the duration of the vegetative lifespan. As such, it is an optimal model system for assessing the impact of life history changes, which modify plant vegetative lifespan, on the evolution of the immunity system. Indeed, in annual (monocarpic) species, which grow and reproduce only once, flowering time marks the end of the vegetative growth phase. Seed production in monocarpic species is terminated by senescence and death, so that flowering time provides a good proxy for lifespan. In *A. thaliana*, it has been scored in a number of conditions (Brachi *et al.*, 2010; Sasaki *et al.*, 2015; Roux & Bergelson, 2016) and flowering time changes are often locally adaptive (Le Corre, 2005; Toomajian *et al.*, 2006; Montesinos-Navarro *et al.*, 2011; Debieu *et al.*, 2013; Li et al., 2014; Hu *et al.*, 2017). Natural variation in flowering time can thus be used to investigate the impact of lifespan changes on host defenses.

The immune system has also been intensively studied in this species, revealing multiple layers of defenses, ranging from basal immunity, which is sufficient to control most microbes, to severe reactions that actively defeat virulent pathogens (Jones & Dangl, 2006). Strain-specific immunity components are likely to be linked in their evolution to the virulence specificity of co-occurring pathogens (de Meaux & Mitchell-Olds, 2003; Moeller & Tiffin, 2005; Roux & Bergelson, 2016). Recent fluctuations in the composition of the pathogen population may therefore affect the specific components of immunity targeted by these epidemics and thereby mask or blur the long-term impact of lifespan modifications. To minimize this effect and to highlight the impact of lifespan variation, we took a genomics approach and examined how flowering time co-varies with expression levels of genes with an experimentally-supported function in immunity. These approximately 700 genes jointly reflect a broad spectrum of traits, which, when their expression increases have a positive effect on immunity (Eulgem, 2005; Vetter *et al.*, 2012; Boccara *et al.*, 2014). We test below whether their expression level, a proxy for their effectiveness, co-variates with flowering time, a proxy for lifespan in the field and further examine the roles played by demographic history and pleiotropy in shaping patterns of co-variation.

## RESULTS

### **Positive co-variation between expression levels of immunity genes and the timing of flowering in Swedish *A. thaliana* populations**

We first focused on a set of 138 genotypes originating from Sweden because high quality data were available for both genome-wide expression profiles and flowering time estimates (Dubin *et al.*, 2015; Sasaki *et al.*, 2015). These two studies were part of a single experiment in which flowering time and gene expression were characterized at both 16°C and 10°C under long day conditions in growth chambers. We focused on the data collected at 16°C and computed Spearman correlation coefficients between the expression level of each gene and flowering time. Of 22,686 genes, for which expression levels could be quantified, 1,374 (6%) were significantly correlated with flowering time under a 5% false discovery rate (FDR). We first verified that genes annotated for their function in flowering time were among the genes whose expression correlates with the phenotype. Overall, genes with an experimentally validated function in flowering time in the genome were not enriched among those genes (6.9% of 630 genes at FDR 0.05, Hypergeometric test, p= 0.19), yet the two well-known regulators of flowering time, FLOWERING LOCUS C and SUPPRESSOR OF OVEREXPRESSION OF CONSTANS-1 (FLC and SOC1, Spearman correlation *ρ*=0.50 and -0.62, FDR-corrected p=2.87e-6 and p=7e-12, respectively) were the two most strongly correlated genes. In addition, using the R-package TopGO, we examined patterns of functional enrichment among genes that tended to be more expressed in early flowering genotypes. Many functional gene ontology (GO) categories related to cell differentiation and growth were enriched (Suppl. Table S1) and the GO category “regulation of flower development” was among the most over-represented (*p*=8.00e-14, Suppl. Table 1). This observation confirmed the biological relevance of the data set examined.

Next, we tested whether immunity genes were enriched among genes whose expression correlated with the timing of flowering. Among genes with significant correlation with the phenotype, we observed a significant excess of immunity genes (8.6% of 691 genes at 5% FDR, hypergeometric test, p=0.002). The distribution of correlation coefficients was also significantly skewed towards higher correlation coefficients for immunity genes (Fig. 2A, Kolmogorov-Smirnov test, p<2.2e-16). GO enrichment analysis showed that genes involved in GO “oxidation-reduction process” and “response to wounding” were among the most strongly enriched (p<1e-30, p=1.1e-19, respectively, Suppl. Table 1). This first analysis revealed a pronounced pattern of positive co-variation between flowering time and the expression of immunity genes.

In laboratory conditions, genotypes with a strong requirement for vernalization tend to show a strong delay in flowering that often does not translate into late flowering in the field (Brachi *et al.*, 2010; Li et al., 2014). Indeed, in the field plants often experience sufficient levels of cold to fulfill their vernalization requirement. In fact, only the 51 genotypes that advanced their flowering time at 16°C compared to 10°C (e.g. those that did not need low temperatures to induce flowering), showed a correlation in their flowering across temperatures (Sasaki *et al.*, 2015). Flowering time variation across the latter sub-sample of genotypes may therefore allow a more accurate classification of genotypes with increasing vegetative lifespan. Among the 507 out of 22,686 (2.2%) genes that displayed a significant positive correlation with flowering time at 10% FDR across this restricted sample of genotypes, 16/630 genes were annotated for their function in flowering. As in the above, several known flowering time regulators were among the genes associated with flowering time, such as FLOWERING LOCUS C (FLC), GIGANTEA, FLOWERING PROMOTING FACTOR 1-LIKE PROTEIN 2 (FLP2) or even the genes PHYTOCHROME INTERACTING FACTOR 4 (PIF4) and PHYTOCHROME INTERACTING FACTOR 5 (PIF5), which had been associated with accelerated flowering (Andrés & Coupland, 2012; Thines *et al.*, 2014). Although the whole set of flowering time genes was not significantly enriched among genes correlating positively with the timing of flowering (2.5%, hypergeometric test, p=0.24), they tended to be more highly expressed in early flowering genotypes (excess of negative correlations, Kolmogorov-Smirnov test, p=1.16e-13, Fig. 2B). The GO category “regulation of flower development” was even more over-represented in this dataset (*p*<1e-30, Suppl. Table 1). Higher expression of genes associated with the positive regulation of flowering was observed among early-flowering genotypes. This further confirms that expression variation was correctly quantified.

We also observed that variation in immunity gene expression tended to correlate positively with variation in flowering time, after excluding vernalization-sensitive genotypes. In total expression of 28 of the 691 genes belonging to the immunity gene category correlated significantly with flowering at 10% FDR. They were significantly enriched (4%, 1.8-fold enrichment, hypergeometric test, p=0.0009). Compared to the ensemble of expressed genes in the genome, they generally tended to be more highly expressed in late flowering genotypes (marked excess of positive correlations, Kolmogorov-Smirnov test, p<2.2e-16, Fig. 2B). GO enrichment analysis showed that genes involved in the GO categories “response to chitin” and “regulation of plant-type hypersensitive response” were the two most strongly enriched (both p<1e-30, Suppl. Table 1). We thus conclude that the correlation between expression of immunity genes and the timing of flowering is independent of allelic variation in vernalization requirements.

### Positive co-variation of immunity gene expression with flowering time is independent of population structure and is also detected in a second sample of broader geographic origin

Relatedness among individuals in the sample may drive the correlation between expression of immunity genes and the timing of flowering. In fact, flowering time in the Swedish lines is strongly associated with the demographic history of these populations and thus with their population structure (Dubin *et al.*, 2015; Sasaki *et al.*, 2015). We therefore also computed for each gene, the correlation between gene expression and flowering time with a mixed-model that included a kinship matrix for the 51 genotypes that lacked vernalization requirement (see methods; Yu *et al.*, 2006; Stich *et al.*, 2008). This analysis revealed that, for immunity genes, the distribution of correlation coefficient estimates remained strongly skewed towards positive values, after population structure was accounted for (Kolmogorov-Smirnov test, p=2.2e-16, Suppl. Fig. 1). However, the whole set of immunity genes was no longer enriched among genes with a significant co-variation with flowering time (5.2%vs. 5%, hypergeometric test, p= 0.6).

**Figure 1:**
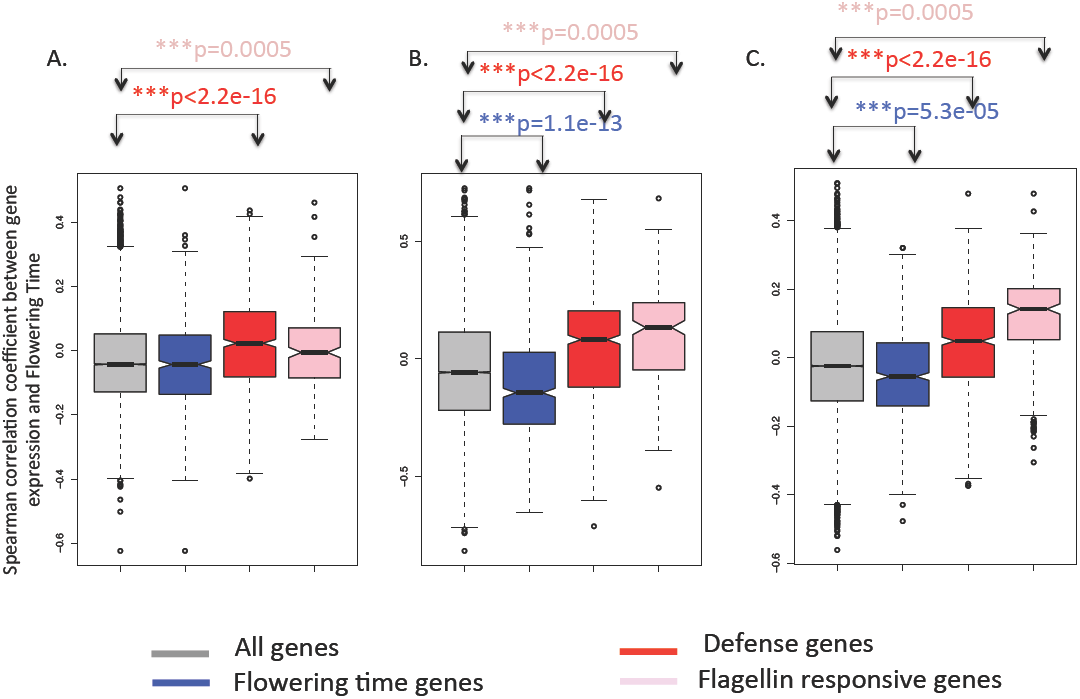
Distribution of Spearman correlation coefficients between expression levels of each expressed *A. thaliana* gene and flowering time. Grey: All expressed genes; Blue: Genes annotated as flowering time genes (FT genes); Red: Genes annotated as immunity genes; Pink: Flagellin-responsive (FlaRe) genes (Navarro *et al.* 2004). **A**. For 138 Swedish genotypes; **B**. Analysis restricted to 51 Swedish genotypes showing correlated flowering time at 10°C and 16°C; C. Species-wide sample of 52 genotypes. Distribution for each group of genes was compared to the genome-wide distribution (black double-head arrow) with a Kolmogorov-Smirnov test. P-values are given in the color corresponding to the gene class. Spearman correlation coefficients were computed between expression levels of each of 23,511 expressed *A. thaliana* genes, reported in Durbin *et al.* 2015 for 9th leaf seedlings, and flowering time measured in the same condition for 51 genotypes originating from natural populations in Sweden (Sasaki *et al.* 2015). *** p < 0.001.

**Figure 2:**
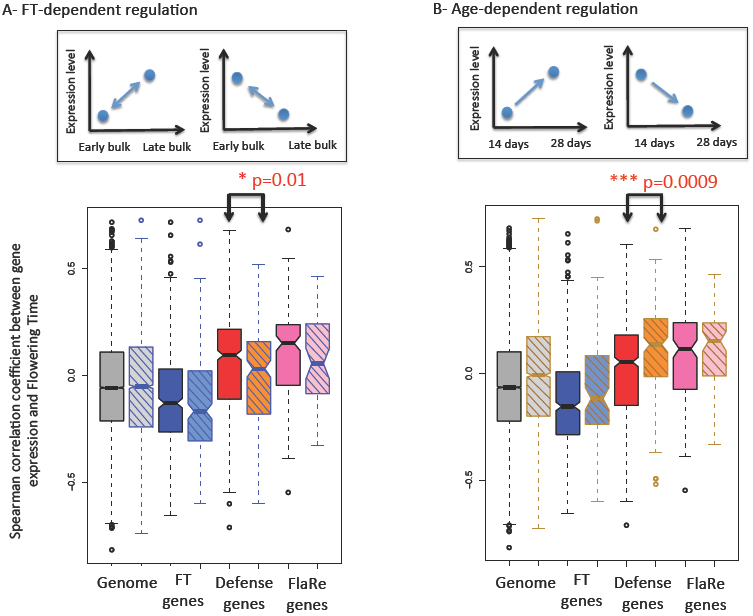
Distribution of Spearman correlation coefficients between gene expression level and flowering time. **A.** Partition of genes controlled by flowering time (hatched boxes with blue border) vs independent from flowering time (uniform boxes with black border); **B.** Partition of genes controlled by development (hatched boxes with orange border) vs independent from development (uniform boxes with black border). Inserts in the top of the figure illustrates how these gene classes were defined. Immunity genes that are not controlled by flowering time but controlled by development tend to have higher correlation coefficients of natural variation for expression with natural variation for flowering time. Grey: All expressed genes; Blue: Genes annotated as flowering time genes (FT genes); Red: Genes annotated as immunity genes; Pink: Flagellin-responsive (FlaRe) genes (Navarro *et al.* 2004). P-values for Kolmogorov-Smirnov test comparing the distribution of genes within each category that are independent of or regulated by **A.** flowering time or **B.** age are shown when significant. Note that only 12 FlaRe genes are controlled by flowering time in our experiment. Spearman correlation coefficients were computed between expression levels of each of 23,511 expressed *A. thaliana* genes, reported in Durbin *et al.* 2015 for 9th leaf seedlings, and flowering time measured in the same condition for 51 genotypes originating from natural populations in Sweden (Sasaki *et al.* 2015). * p < 0.05, *** p < 0.001.

We note that accounting for population structure also did not change the pattern of co-variation between gene expression of flowering genes and timing of flowering itself. They showed a coefficient distribution that was strongly skewed towards negative values (Kolmogorov-Smirnov test, p=2.2e-16) and were significantly over-represented among genes with expression significantly associated with flowering time (8% vs 5% at 5% FDR, hypergeometric test, p=0.0005).

We further investigated whether the skew towards positive co-variation between immunity gene expression and flowering time is limited to the regional subset of genotypes growing in Sweden or whether it is a feature of diversity that segregates across the whole range of the species. For this, we turned to a species-wide dataset of gene expression variation collected in young seedlings (Schmitz *et al.*, 2013). For 52 of these genotypes, the duration of vegetative growth had been determined under natural conditions in the field (Brachi *et al.*, 2010). Although a skew towards negative correlation for flowering time genes was observed (Kolmogorov Smirnov test, p=5.3e-5, Fig. 2C), the seedling of these earlier flowering genotypes did not yet express genes important for the formation of flower (Suppl. Table 1).

Nevertheless, we again observed a strong skew towards positive correlation between immunity gene expression and flowering time, indicating that genotypes that will flower later expressed them at a higher level (Kolmogorov Smirnov test, p<2.2e-16, Fig. 2C). Immunity genes were not particularly enriched among genes with significantly correlated expression and flowering time at 5% FDR (5% for both, hypergeometric test, p=0.24). Yet, GO categories such as “response to chitin”, “respiratory burst involved in immunity response”, “response to wounding” and immunity response to fungus” were the four most strongly enriched functions among genes with highest Spearman correlation coefficients (all *p*<1e-30, Suppl. Table 1).

Contrasting genotypes of diverse flowering time (e.g. lifespan) revealed that, in natural populations, immunity genes tend to co-vary positively with this trait. The latter two analyses showed that this effect remained when population structure was accounted for and was also detectable in another gene expression dataset and with a different set of genotypes.

### A bulk-segregant analysis demonstrates that co-variation is not due to pleiotropic effect of flowering time control

The tendency of immunity genes to show expression levels correlating positively with flowering may be due to the pleiotropic action of regulatory genes that co-regulate flowering time and immunity. In plants, the impact of development and growth regulators on defense systems is being increasingly recognized (Alcázar *et al.*, 2011). There is evidence that flowering time and defense control each other (Korves & Bergelson, 2003; Pagán *et al.*, 2008; Fan *et al.*, 2014; Lozano-Durán & Zipfel, 2015; Jiménez-Góngora *et al.*, 2015; Davila Olivas *et al.*, 2017; Develey-Riviere & Galiana, 2007; Pajerowska-Mukhtar *et al.*, 2009; Martinez *et al.*, 2004; Whalen, 2005; Kerwin *et al.*, 2015; Lyons *et al*. 2015). If so, the pattern we observed would not reflect the joint optimization of immunity and life history strategy but only the pleiotropic action of their regulators. We therefore asked to which extent flowering time and the expression of immune-related genes could be genetically separated and thus evolve independently.

We therefore designed an experiment to describe the level of pleiotropy of flowering time regulators on the expression of immunity genes. If such regulators control the pattern of covariation reported in Fig. 2, it should not be possible to separate variation in immunity gene expression from variation in flowering time in a segregating recombinant inbred population. We used the two genotypes Col-0 and Bur-0, which differ in flowering time (Simon *et al.*, 2008) and were also reported to exhibit markedly distinct sensitivities to flagellin, with the later flowering genotype Bur-0 displaying stronger basal immunity (Vetter *et al.*, 2012). We analyzed the transcriptomes of these two lines at 14 and 28 days after germination (see methods) and found that the skews shown in Fig. 2 remain when the dataset was reduced to the genes that differed in expression between these two lines (Suppl. Fig. 2). This confirmed that these two genotypes could help identify immunity genes that share genetic regulators with flowering time.

We designed a cost-effective approach to identify the genes whose expression variation cannot be separated from flowering time. We used 244 recombinant inbred lines (RILs) derived from a cross between the parents Bur-0 and Col-0, followed by >8 generations of selfing (Simon *et al.*, 2008). We bulked RILs by their flowering time and characterized their transcriptomes at 14 and 28 days after germinations using RNA sequencing (see methods). In RILs, the genomes of the parental genotypes are randomly shuffled by recombination. Because of this genetic property, RILs are commonly used to identify Quantitative Trait Loci (QTL), which are genomic regions underlying the genetic control of phenotypic variation. In our approach, this means that differences in gene expression between early- and late-flowering RILs reflect differences that are genetically associated with flowering time. The experimental strategy is described in Suppl. Fig. 3-4. This strategy does not allow characterizing the exact genetic architecture of gene expression variation, but it allows the identification of genes whose expression variation is controlled either by flowering-time regulators or by genes located in the genomic vicinity of these regulators. Thereafter, we named these genes <u>F</u>lowering-<u>T</u>ime (FT)-dependent genes.

**Figure 3:**
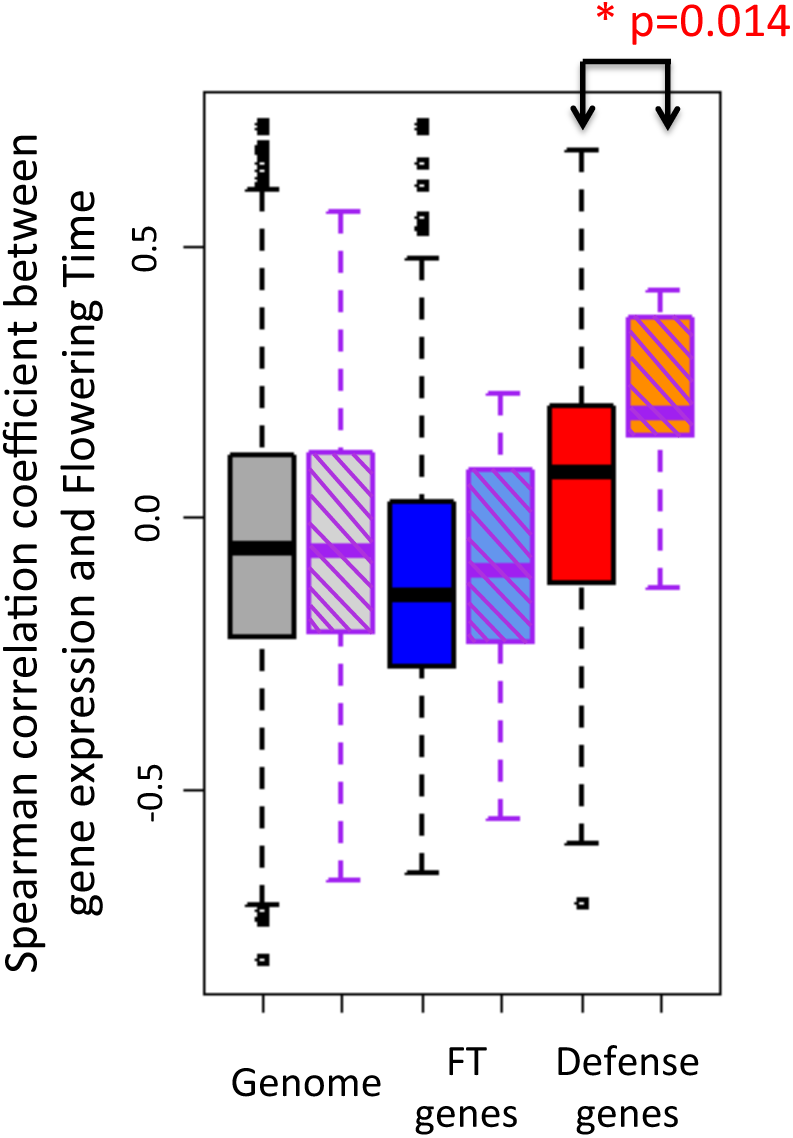
Distribution of Spearman correlation coefficients between gene expression level and flowering time. All expressed genes–uniform boxes with black border-vs. genes with fitness-associated SNPs in Fournier-Level *et al.* (2011), - hatched boxes with purple border-. Grey: All expressed genes; Blue: Genes annotated as flowering time genes (FT genes); Red: Genes annotated as immunity genes. Immunity genes that carry SNPs associating with fitness tend to have higher correlation coefficients of natural variation for expression with natural variation for flowering time. P-values for Kolmogorov-Smirnov test comparing the distribution for genes within each category are shown when significant. Spearman correlation coefficients were computed between expression levels of each of 23,511 expressed *A. thaliana* genes, reported in Durbin *et al.* 2015 for 9th leaf seedlings, and flowering time measured in the same condition for 51 genotypes originating from natural populations in Sweden (Sasaki *et al.* 2015). * p < 0.05.

Of a total of 20,553 genes expressed in both the parental genotypes and RIL pools, 6,097 (29%) were differentially expressed between early and late flowering RIL pools, i.e. FT-dependent. As expected, there was a strong excess of genes annotated as having a function in flowering time among FT-dependent genes (223/630 – 36%, hypergeometric test, p=3.7e-5). This demonstrated that this strategy effectively highlighted genes whose expression is under the genetic control of flowering time regulators. By contrast, immunity genes were not over-represented among FT-dependent genes. More so, they were clearly under-represented among FT-dependent genes at the second time point of sampling (1.15 fold less abundant than expected by chance at day 28, hypergeometric test, p=0.01). Only 19% of all immunity genes were FT-dependent. These genes, however, did not explain the skew towards positive co-variation with flowering time reported in Fig. 2. Immunity genes, whose expression was not differently expressed between RIL pools (i.e. genes whose expression is not dependent on the regulators of flowering time), in fact, tended to be more skewed towards positive correlation coefficients than FT-dependent immunity genes (Kolmogorov-Smirnov test, p=0.01, Fig. 2A). We observed that FT-dependent flowering time genes did not shift significantly from the distribution of correlation in the rest of the genome (Kolmogorov-Smirnov test, p=0.15, Fig. 2A). Therefore, the excess of positive expression co-variation with flowering time observed among immunity genes is most strongly driven by genes whose expression level was easily separated from variation in flowering by recombination.

### Age-regulated immunity genes often show positive co-variation with flowering time

Immunity genes are often observed to change their activity with age and development (Barton & Boege, 2017). Because we had sampled material at day 14 and day 28 after germination, we could also separate genes whose expression changed with age (here after named age-regulated genes) from genes with similar expression levels in 14- and 28-day-old plants (see methods). Age-regulated genes were markedly more frequent among annotated immunity genes than among annotated flowering time genes (243/630 – 38% vs 334/691 - 48%, for flowering-time and immunity genes, respectively, Chi Square test, p= 7.2e-11). In *A. thaliana,* a so-called age-related resistance is activated in older *A. thaliana* plants, providing them with a immunity barrier against a broad spectrum of pathogens (Rusterucci *et al.*, 2005). In agreement with our findings, the timing of age-related resistance had been reported not to stand under the direct control of flowering time (Wilson *et al.*, 2013).

The subset of genes, whose expression variation in natural populations correlated with flowering time, were also enriched among age-regulated genes (hypergeometric test, p= 7.2e-11). Altogether, 4% (348/8565) and 6% (498/7935) of age-independent and age-regulated genes, respectively, were correlated with flowering time at 5% FDR. Immunity genes contributed significantly to this excess, because the expression levels of immunity genes that were age-regulated tended to show a strong skew towards positive correlation with flowering time in natural populations (Fig. 2B, Kolmogorov-Smirnov test, p=0.0009). Our analysis thus indicates that the tendency of immunity genes to co-vary positively with flowering time in natural population is i) not explained by the genetic control of flowering time and ii) increased among genes whose expression is regulated by plant age.

### Genes activated by elicitors of basal immunity also show an excess of positive correlations with flowering time

In the above analyses, immunity levels were represented by a set of 731 genes annotated for functions related to immunity. To test whether this trend towards positive covariation between immunity gene expression and flowering time was limited to the set of genes defined by Gene Ontology categories, we analyzed an independent set of immunity-related genes: the 245 genes whose expression is activated in *Arabidopsis* seedlings upon perception of flagellin by the PAMP receptor kinase FLAGELLIN SENSING 2 (FLS2), hereafter named FlaRe genes (Navarro *et al.*, 2004). FlaRe genes coordinate cellular and developmental responses to exposure of molecular signatures of bacteria. Only 10 FlaRe genes overlapped with the immunity-annotated genes used above. We observed that FlaRe genes were enriched among genes showing positive co-variation with flowering time (Fig. 2A-C, Kolmogorov-Smirnov test, p< 2.2e-16). This observation remained when accounting for population structure (Suppl. Fig. 1, Kolmogorov-Smirnov test, p<2.2e-16) and was also seen for flowering time measured in the field in a species-wide sample of genotypes (Fig. 2C, Kolmogorov-Smirnov test, p<2.2e-16). When partitioning genes according to whether or not they were FT-dependent or age-regulated, we observed that FT-dependence did not significantly change the distribution of correlation coefficients between FlaRe gene expression and flowering time across natural genotypes (Fig. 2A-B, Kolmogorov-Smirnov test, p=0.15 and p=0.32, for FT-controlled and age-regulated genes, respectively). Nevertheless, FlaRe genes were significantly under-represented among FT-dependent genes, especially at the second sampling time point (1.8-fold less frequent among flowering time controlled genes, hypergeometric test, p= 2.24e-05). By contrast, they were over-represented among age-regulated genes (2.1-fold more frequent among age-regulated genes, hypergeometric test, p= 2.6e-08). Thus, the positive co-variation reported in Fig. 1A-C is unlikely to result from the pleiotropic action of flowering time regulators on FlaRe genes. This suggests that, like for annotated immunity genes, alleles attenuating the expression of FlaRe genes were assorted with early-flowering alleles in natural populations and vice versa.

### Fitness-associated immunity genes show higher correlation coefficients with flowering time

We further asked whether genes with fitness-relevant variation have expression levels that are more strongly assorted with variation in the timing of flowering. A reciprocal transplant experiment performed in 4 locations throughout Europe identified 866 nucleotide variants in the genome of *A. thaliana* that significantly associated with fitness differences manifested in natural conditions (Fournier-Level *et al.* 2011). Of these variants, 15 mapped to immunity genes and 17 to flowering genes. Association with fitness coincided with a skew towards higher correlation coefficients for immunity genes only (Fig. 3, Kolmogorov-Smirnov test, D=0.46, *p*=0.014 and p>0.05 for immunity and flowering time genes, respectively). One of the immunity genes (AT3G16720), which is activated upon exposure to the fungal PAMP chitin, was FT-dependent but it did not explain this pattern (Kolmogorov-Smirnov test, *p*=0.028 without AT3G16720). Five of the immunity genes with FT-independent immune functions were age-regulated (AT1G18150, AT1G80840, AT4G01700, AT5G19510, AT5G57220) but this did not explain the pattern either (Kolmogorov-Smirnov test, *p*=0.009 without these genes). Of the 245 FlaRe genes, 3 contained fitness-associated SNPs. These three genes were among the genes with highest correlation coefficients (AT1G19670: ρ=0.397, AT3G16720: ρ=0.282, AT4G38860: ρ=0.487). We thus observe that immunity genes that can be most relevant for fitness in natural populations of *A. thaliana* are also genes whose expression levels were most strongly assorted with alleles determining flowering time.

## DISCUSSION

### **Evidence for concerted evolution of immunity and flowering time in *A. thaliana***

Our analyses reveal that, in *A. thaliana*, individuals with a shorter vegetative lifespan tend to express immunity genes at a lower level. The bulk analysis of early- and late-flowering RILs shows that this pattern of co-variation results from the combination of independent alleles controlling immunity gene expression and flowering time in natural populations, because these alleles could be separated in the segregating recombinant offspring of an early- and a late-flowering genotype. Because co-variation is also i) robust to the demographic history of the populations and ii) particularly pronounced for immunity-gene variants that associate with fitness, our analyses suggest that this allelic combination is assembled by natural selection. This pattern is confirmed by the examination of genes annotated with a function in immunity and genes observed to respond to elicitation by the common bacterial elicitor flagellin. Our data further suggests that much of the positive co-variation between immunity gene expression and flowering depends on plant age. This factor is of recognized importance in plant immunity (Alcázar *et al.*, 2011; Lozano-Durán & Zipfel, 2015; Carella *et al.*, 2015) and also very well documented in ecological studies (Barton & Boege, 2017). Based on our findings, it is tempting to speculate that variation in age-dependent regulation of immunity may mediate the co-variation we report.

### Co-variation between immunity and flowering time is not explained by variation in vernalization requirements

Flowering time variation depends on seasonal fluctuations, on the timing of germination and on the genetics of its control (Lempe *et al.*, 2005; Balasubramanian *et al.*, 2006; Korves *et al.*, 2007; Burghardt *et al.*, 2015; Hu *et al.*, 2017). Genotypes with a strong vernalization requirement, which are thought to have an obligate winter annual strategy, contribute strongly to the variation reported in the literature because they display much delayed flowering in the laboratory (Lempe *et al.*, 2005; Li et al., 2014; Sasaki *et al.*, 2015). The pattern we report, however, is not due to the assortment of immunity gene expression variants with alleles imposing a strong vernalization requirement. Indeed, since we observed that the pattern of co-variation between immunity gene expression and flowering time is magnified in plants whose flowering is not accelerated by cold exposure (Fig. 2A-B), this pattern is not driven by the genotypes requiring vernalization. In addition, the same pattern of co-variation is observed in a global sample of ecotypes, whose flowering time was scored in an outdoor common garden experiment, where plants were naturally vernalized (Fig. 2C). Therefore, we believe that the flowering time measures we used here do capture some of the natural lifespan variation. Future studies will have to confirm that flowering time variation scales with average differences in the lifespan expressed at the location of origin of each genotype.

### Positive co-variation between lifespan and immunity suggests cascading effect of flowering time adaptation on immunity evolution

Two alternative scenarios may lead to concerted evolution of flowering time and immunity. First, in conditions where disease pressure is high, both shorter lifespan and stronger immunity can be expected to be advantageous, in order to simultaneously minimize the probability of attack, and maximize the probability of survival in case of attack. Under such scenario, negative co-variation between immunity and lifespan is expected. Alternatively, if lifespan is evolving under evolutionary forces independent of disease pressure, a reduced probability to encounter pathogens will favor mutations transferring energy allocated to immunity into energy allocated to growth. Indeed, defensive functions are known to be costly for the organism (Lochmiller & Deerenberg, 2000; Purrington, 2000). As a consequence the allocation into immunity is predicted to decrease where shorter lifespan evolves. Under this second scenario, a pattern of positive co-variation is expected between immunity and lifespan.

The pattern of co-variation we report here for immunity vs. flowering time is indeed positive and thus lends support to the second scenario. Local adaptation of flowering time is well documented in *A. thaliana* (Le Corre, 2005; Toomajian *et al.*, 2006; Méndez-Vigo *et al.*, 2011; Brachi *et al.*, 2013; Li et al., 2014; Debieu *et al.*, 2013; Burghardt *et al.*, 2015; Vidigal *et al.*, 2016). In addition, several studies support the idea that increased basal level in immunity components improves immunity (Vetter *et al.*, 2012; Boccara *et al.*, 2014). At the same time, variants involved in the surveillance systems directed against pathogenic virulence factors were shown to incur substantial fitness costs (Tian *et al.*, 2003; but see also MacQueen *et al.*, 2016) and variation in basal immunity was negatively correlated with plant growth (Vetter *et al.*, 2012). Our results are thus compatible with an evolutionary scenario in which local adaptation of flowering time has cascading effect on immunity, possibly because a reduction of the plant’s lifespan increases the cost/benefit ratio of immunity. This may also explain why genes involved in local adaptation in China are enriched among both flowering time and immunity genes (Zou *et al.* 2017).

However, a positive pattern of covariation could also arise even if the two traits evolve independently. Indeed, it is possible that populations where early flowering is advantageous coincide with populations where disease pressure is lower and *vice versa*. We cannot formally exclude that this scenario does not apply, because too little is known about variation in disease pressure in *A. thaliana* natural populations. Several elements, however, indicate it is unlikely. First, the rapid cycling genotypes are more frequent at intermediate latitudes, where summers are mild and wet (Lempe *et al.*, 2005; Debieu *et al.*, 2013). Since these conditions are also favorable to diseases, it is unlikely that higher disease pressure is found in areas where delayed flowering is more adaptive. Second, it is unlikely that this pattern may be due to herbivore enemies. Indeed, more severe herbivory damage has been observed on early-flowering *A. thaliana* individuals grown in the field (Weinig *et al.*, 2003). This seems to be common in plant species and should select for higher defense among early-flowering genotypes (Carmona *et al.*, 2010). Third, such scenario would assume that variation in disease pressure does not alter the trade-off between survival and reproductive output. This trade-off, however, is central in many models explaining the evolution of the timing of flowering in monocarpic plant species (Mitchell-Olds, 1996; Metcalf & Mitchell-Olds, 2009; Ashworth *et al.*, 2016).

Our results are therefore compatible with a scenario, in which adaptation of life history traits has a cascading effect on the evolution of immunity in *A. thaliana*. These findings do not contradict evidence that a tug of war characterizes the evolution of pathogen-specific components of immunity (Tellier & Brown, 2007; Roux & Bergelson, 2016). Indeed, by examining the basal expression level of a large set of genes involved in the immune reaction, the impact of durable selective forces on general immunity levels can be detected. This approach circumvents the potentially confounding signature left by a recent epidemics on strain-specific R-genes. Indeed, testing phenotypic variation in disease resistance across genotypes with different life-history alleles would probably reveal variation in gene-for-gene resistance, but the pervasive impact of selection fine-tuning energetic costs associated with immunity strategies would remained masked.

Interspecific differences in the investment in defence against herbivory has been often associated with differences in lifespan and growth rate (Endara & Coley, 2010; Kooyers *et al.*, 2017). Future studies will also have to examine whether a similar evolutionary trend has emerged in species that have reshaped their life history to decrease overall vegetative lifespan. Early flowering is actually often favored when the favorable season is shortened (Franks *et al.*, 2007; Kenney *et al.*, 2014). Ongoing selection for early flowering is clearly widespread at temperate latitudes (Munguía-Rosas *et al.*, 2011) and transitions from perenniality to annuality occur frequently within phylogenies (Kiefer *et al.*, 2017). Testing whether life span reduction associates with an attenuation of immunity gene expression should therefore be possible in many taxa.

### The impact of life history evolution on defense systems is expected across all kingdoms

In animals, the idea that the optimal investment in immunity depends on the life history of a species was also incorporated in evolutionary models (Jokela *et al.*, 2000). For plants and animals alike, resources available to the organism are limited. Energetic demands on growth may compete with those required for mounting immunity or counteracting the negative effects of parasites and pathogens (van Boven & Weissing, 2004; Lazzaro & Little, 2009; Dowling & Simmons, 2009; Seppälä, 2015). Several evolutionary models show that a prolonged lifespan is predicted to favor resource investment into immunity (Jokela *et al.*, 2000; Medzhitov & Janeway, 2000; van Boven & Weissing, 2004; Miller *et al.*, 2007). As a consequence, changes in life history can mold the evolution of immune systems in animals as well (Van Valen, 1973; Sheldon & Verhulst, 1996; Schulenburg *et al.*, 2009). This theoretical prediction is supported by analyses of sexual dimorphism in the duration of effective breeding: females with increased reproductive longevity show stronger immune-competence but also by a meta-analysis of selection experiments (Rolff, 2007; Nunn *et al.*, 2009, van der Most *et al.* 2011). In frogs, fast developing species were also shown to be more susceptible to infection by trematodes (Johnson *et al.*, 2012). Yet, such studies cannot exclude that longevity and immunity are constrained in their evolution by common regulatory factors or causal inter-dependence. To the best of our knowledge, this study is the first to provide evidence that natural variation in the activity of genes that are important for defeating pathogens is assorted with alleles controlling variation in a life history trait of considerable importance for adaptation. Local adaptation for lifespan should therefore be considered as a potentially important contributor to the maintenance of genetic diversity in immune systems.

## MATERIAL AND METHODS

### Flowering and immunity candidate genes

Gene Ontology (GO) categories were used to identify functionally related genes whose annotation was inferred from experiments, direct assays, physical interaction, mutant phenotype, genetic interactions or from expression patterns. Based on the keyword “flowering” in the TAIR database, 659 flowering time genes were selected. For immunity genes, we united 17 GO categories yielding 731 genes (Suppl. Table S2). For flagellin responsive (FlaRe) genes, we took the set of 245 genes that were activated in seedlings described in (Navarro *et al.*, 2004) (Suppl. Table S2). Subsets of flowering, immunity and FlaRe genes containing fitness-associated single nucleotide polymorphisms (SNPs) were retrieved from Fournier-Level *et al.* 2011.

### Correlation between gene expression and flowering time in a natural population

We analyzed two published sets of natural ecotypes for which both genome-wide expression profiles and flowering time estimates were available. The first dataset comprised 138 lines from Sweden scored for both flowering time (for plants grown at 16h light-8hour dark at constant 16°C) and gene expression in whole rosette collected at the 9-true-leaf stage (Dubin *et al.*, 2015; Sasaki *et al.*, 2015). For this first dataset, gene expression and flowering were determined in the same experiment. The second dataset combined data from two sources. RNA extracted from 7-day old seedlings of 144 genotypes grown on agar plate in long days had been sequenced (Schmitz *et al.*, 2013) and expression levels quantified as quantile normalized fragment numbers per kilobases and million reads (FPKM). For 52 of these genotypes, flowering time, measured in cumulative photothermal units, had been scored in the field (Brachi *et al.*, 2010). Photo-thermal units sum up the combination of temperature and day length and thus provide an estimate of the duration of the favorable season.

Expression counts were loge +1-transformed to include null values of expression and a Spearman correlation coefficient between flowering time and expression level was computed for each gene. P-values were adjusted for false discovery rate using the p.adjust function in R (Benjamini & Hochberg, 1995; Yekutieli & Benjamini, 1999). A Kolmogorov-Smirnov test was used to compare the distribution of Spearman correlation coefficients ρ of flowering time and immunity genes with the distribution of ρ for 22,686 genes for which gene expression was quantified. Gene enrichments were tested using hypergeometric tests in R. The GO enrichment analysis was performed with the Gene Set Enrichment Analysis (GSEA) test akin to non-parametric Kolmogorov-Smirnov tests, first described by Subramanian *et al.*, 2005, and implemented in the “topGO” R package (Alexa and Rahnenfuhrer, 2010). We further applied the *elim* procedure, available in this package, which calculates enrichment significance of parent nodes after eliminating genes of significant children nodes. This controls for the dependency among nested parent-child GO categories so that the significance of each enrichment can be interpreted without over-conservative p-value corrections for multiple-testing (Alexa et al. 2006). To test the impact of population structure on the correlation, we ran a mixed model with the help of the R package *lmekin.* For each gene, we used gene expression level as a dependent variable. Flowering time was used as independent variable and a kinship matrix, generated with a matrix of SNPs segregating among Swedish genotypes (Dubin *et al.*, 2015), was included as random effect. The estimate of the flowering time effect was extracted. This allowed compared the distribution of estimates observed for the whole genome, the subset of flowering time genes, or the subsets of defense genes.

### Analysis of gene expression in segregant pools bulked by flowering time

Seeds of Bur-0, Col-0 and 278 Bur-0xCol-0 Recombinant Inbred Lines (RIL) obtained after 8 generations of selfing were provided by the Arabidopsis Stock Center at INRA Versailles (France, (Simon *et al.*, 2008). Each line was grown individually in six replicates, each in 6cm diameter pots randomly allocated to 24 trays, each containing 35 pots. Seeds were stratified at 5°C for 3 days and grown in growth chambers (Elbanton BV, Holland, equipped with Sylvania Gro-Lux F36W /Gro (T8) fluorescent tubes and Osram 25 W 220 Lumen light bulbs) under long-day conditions (21°C, 16h light, 18°C, 8h dark). Trays were rotated within the chamber every other day. Flowering time was scored as the day to the first open flower. Genotypes of individuals lines were retrieved from Simon *et al.* (2008) and mapping of flowering time recovered the same QTL (not shown).

We selected the 40 RIL in the 15% and 85% quantiles of flowering time for RNA sequencing. Each RIL and the two parental lines were planted in 20 replicates in the conditions described above. At days 14 and 28, the oldest true leaf was flash-frozen in liquid nitrogen. Three pools, each combining 13 RIL, were produced at each time point for early and late lines, for a total of 3 biological replicates, 2 pool types (early and late RIL) and 2 time points (14 and 28 days). For each of the two parental lines, leaves of 12 replicates were pooled for each time point.

RNA was isolated using the TRIzol extraction protocol (ThermoFisher Scientific, USA). DNA traces were removed with the Ambion DNA-free kit (ThermoFisher Scientific, USA) and purified RNA was stored in TE buffer at -80°C. RNA quality and integrity was confirmed with the 2100 Expert Software on a Bioanalyzer (Agilent Technologies, Inc. Waldbronn, Germany). All samples had RNA integrity index (RIN) larger than 8. Single-read libraries were prepared with 1μg of total RNA per sample using the Illumina TruSeq RNA Sample Preparation Kit v2 (Illumina Inc. San Diego, USA) based on poly-A RNA purification. Sequencing of 75bp single reads was performed on the Illumina HighScan SQ system of the Core Facility of the Department of Genetic Epidemiology, Institute of Human Genetics, University of Münster, Germany. Raw data has been deposited in NCBI’s Gene Expression Omnibus (Edgar *et al.*, 2002) and are accessible through GEO Series accession number GSE97664.

### Data analysis of RNA-seq from bulk segregant pools

In total, 24 RNA libraries were sequenced. Raw sequences were demultiplexed and read quality validated with FastQC. Bad quality base calls were trimmed using the fastx-toolkit (Version 0.013, Li *et al.* 2009). Trimmed reads (FastQ, quality score 33, quality threshold 20 and minimum length 30 base pair) were mapped to the *A. thaliana* TAIR10 annotated transcriptome using Bowtie 2 (version 2-2.0.0-beta6, (Langmead & Salzberg, 2012). Tophat (version-2.0.5.Linux_x86_64) was used to discover splice sites and Cufflinks for assembling the transcriptome (Trapnell *et al.*, 2010). In total 411,5 M sequence reads were obtained, with a mean read count per sample of 17,1 M reads. After trimming, 96.5% of the reads were mapped uniquely with a final average coverage of 66 reads per base pair.

We used a custom R script to verify that coverage was uniform across transcripts and confirmed that the RNA sequenced was not degraded. Read counts were calculated by counting the number of reads that mapped uniquely to the corresponding gene (isoforms were not considered). Lowly expressed genes with less than 20 reads over all samples were excluded from the analysis. The samples clustered by time point of sampling (Fig. 2), with the exception of RNA samples from the Col-0 at 28 days, which resembled more expression levels measured at 14 days, probably because of its early shift to flowering. Differentially expressed (DE) genes were identified by running a nested analysis of sampling time effects within parental genotype (and/or early- and late-flowering leaf pools) with DESeq2 version 1.2.5 (Anders *et al.*, 2013; Love *et al.*, 2014). P-values were corrected for false discovery rate (Benjamini-Hochberg correction; (Benjamini & Hochberg, 1995). DE genes were defined as having an adjusted p-value<0.05. This analysis allowed the identification of genes showing differential expression between the parents (Suppl. Table S3) and genes showing flowering time dependent expression (differential expression between early and late flowering RIL pools, i.e. FT-regulated genes Suppl. Table S4) both at day 14 and at day 28. We performed further analyses to disentangle significant sources of gene expression variation. To test whether gene expression was significantly modified at each time point, separate tests were performed for each parental genotype and RIL pool type. Genes differentially regulated at 14- and 28-days in Bur-0 (adjusted p-value<0.05) were defined as age-regulated genes (Suppl. Table S5). To determine whether one or both sampling time points drove significant differential expression, separate tests were performed for each time point (not shown).

### Confirmation with qRT PCR

We confirmed gene expression levels for 11 selected immunity genes with differential expression between Bur-0 and Col-0 or early vs. late flowering pools (log2-fold change > 1.5) using RT-PCR. We followed standard protocols and used RNA Helicase (AT1G58060), Protein Phosphatase 2A Subunit A3 (PP2AA3) and transcript AT5G12240 as control genes. Gene expression based on RNA sequencing and RT-PCR were strongly correlated (Pearson correlation, 0.58<R<0.96, max p <0.01).

## Supplementary Tables

**Suppl. Table 1:** GO categories enriched among genes correlating either positively or negatively with flowering time.

**Suppl. Table 2:** List of immunity genes and GO categories (only gene annotations based on experimentally validated open reading frames were considered). FlaRe genes and flowering time genes used in the study.

**Suppl. Table 3:** FT-dependent genes (DE between early and late flowering RIL pools). Output of gene expression analysis includes mean read count (FPKM), log2 fold-change, Standard error of the log2 fold-change; p-value and FDR adjusted p-value. FT-dependent genes have FDR adjusted p-values <0.05.

**Suppl. Table 4:** Differentially expressed genes between Col-0 and Bur-0. Output of gene expression analysis includes mean read count (FPKM), log2 fold-change, Standard error of the log2 fold-change; p-value and FDR adjusted p-value. Genes differently regulated between Col-0 and Bur-0 have FDR adjusted p-values <0.05.

**Suppl. Table 5:** Age-regulated genes defined as differential gene expression changes in Bur-0 between 14- and 28-days. Output of gene expression analysis includes mean read count (FPKM), log2 fold-change, Standard error of the log2 fold-change; p-value and FDR adjusted p-value. FT-dependent genes have FDR adjusted p-values <0.05.

## Supplementary Figures

**Suppl. Figure 1:**
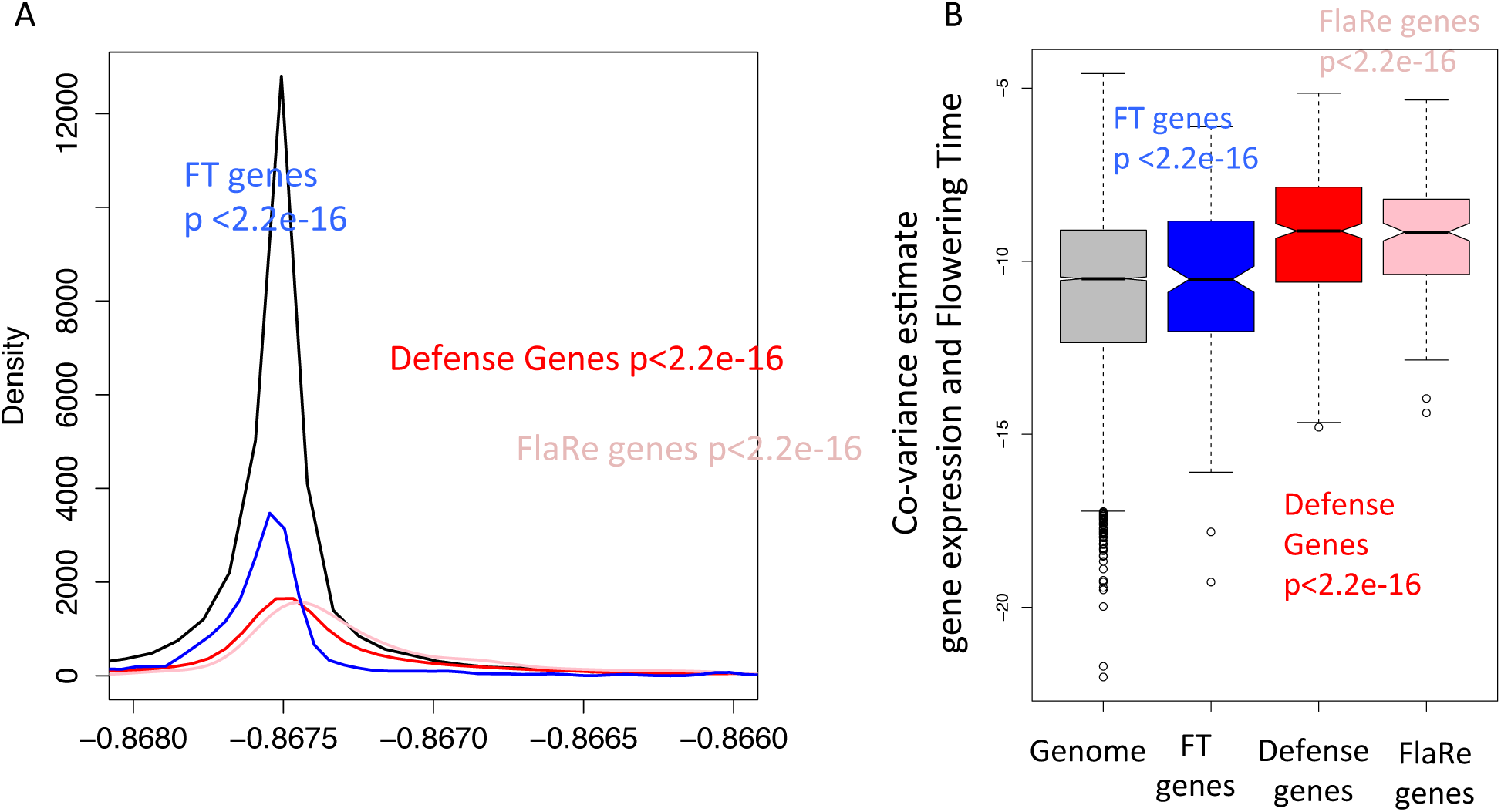
Distribution of estimated effect of flowering time as explanatory factor of gene expression variation, taking into account population structure between Swedish genotypes of group A (Group A genotypes advance their flowering at 16°C compared to 10°C and show correlated flowering at 10°C and 16°C, Sasaki *et al.* 2015). The trend shown in Figure 1 is maintained after accounting for population structure. P-values for Kolmogorov-Smirnov test comparing estimate distribution for the gene subset compared to the genome-wide distribution are given. **A.** Density distribution, **B.** Boxplots.

**Suppl. Figure 2:**
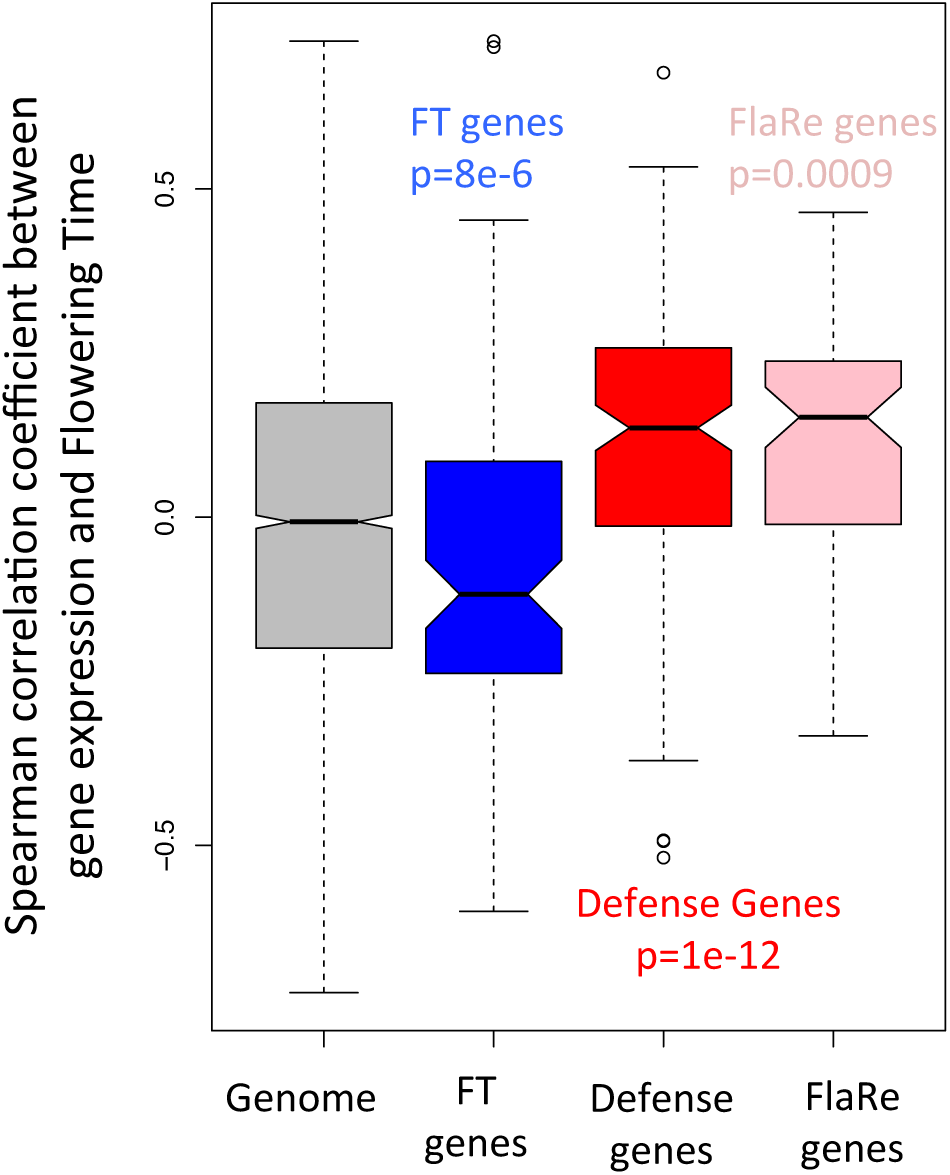
Distribution of correlation coefficients between gene expression and flowering time restricted to the genes showing differential expression in Col-0 vs. Bur-0. Expression differences between the genotypes Col-0 and Bur-0 recapitulate the pattern reported within natural populations in Figure 1. Distribution of Spearman correlation coefficients between gene expression level and flowering time for the set of genotypes showing consistent differences in flowering at 10°C and 16°C (Sasaki *et al.* 2015). This analysis is restricted to the 6980 genes showing differential expression between Col-0 and Bur-0. Immunity genes have a stronger skew in correlation coefficients with flowering time. Spearman correlation coefficients were computed between expression level of each of 6980 expressed *A. thaliana* genes, reported in Durbin *et al.* 2015 for 9th leaf seedlings, and flowering time measured in the same condition. Genotypes originate from natural populations in Sweden (Sasaki *et al.* 2015). Black line: All expressed genes, Blue lines: Gene annotated as flowering time genes (FT genes), Red lines: Genes annotated as immunity genes, Pink line: Flagellin-responsive (FlaRe) genes (Navarro *et al.* 2004).

**Suppl. Figure 3:**
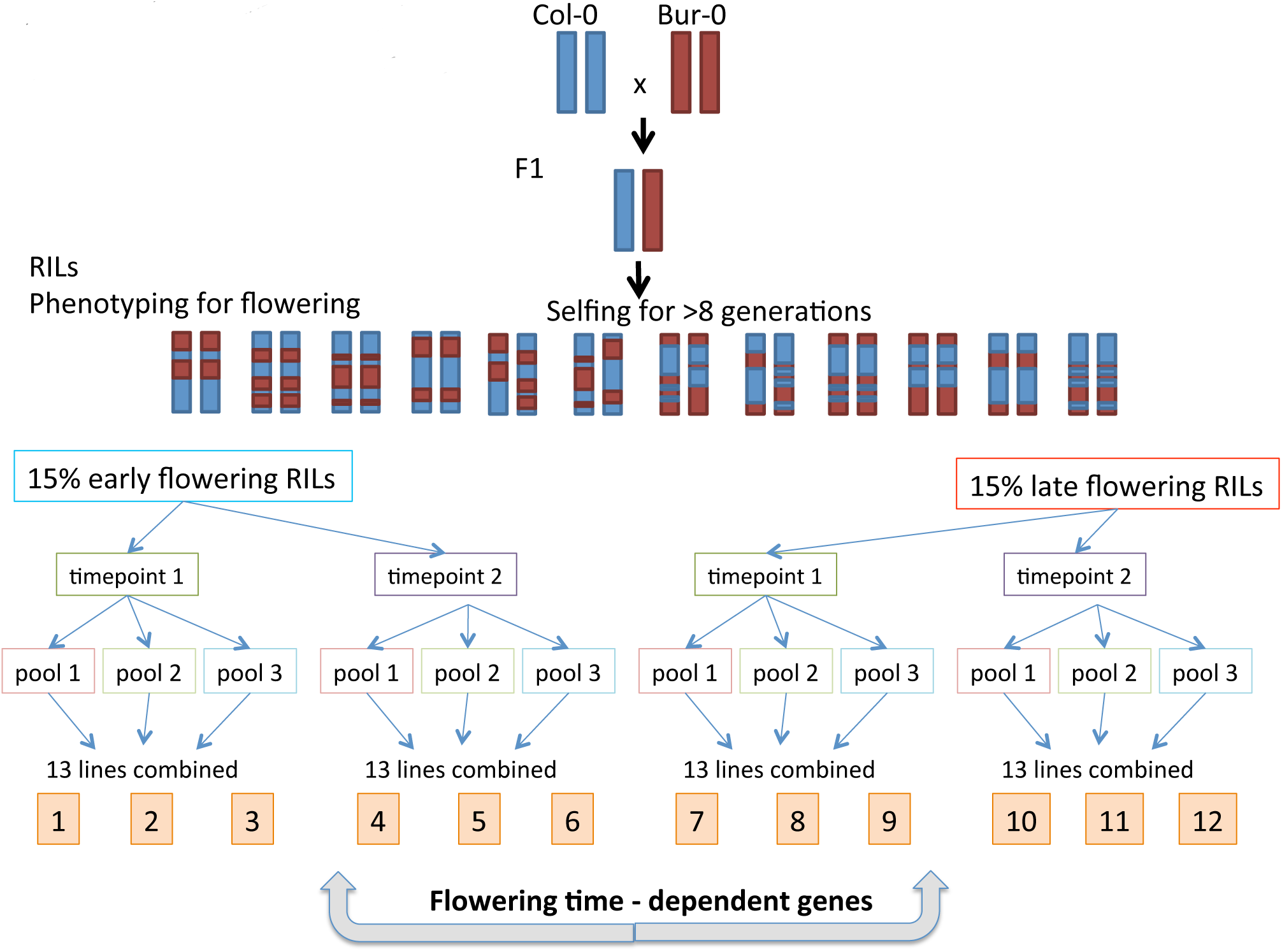
Bulk-sequencing strategy used to identify FT-dependent genes, i.e. genes whose expression is genetically controlled by flowering time regulators or by genes located closely to and therefore co-segregating with flowering time regulators.

**Suppl. Figure 4:**
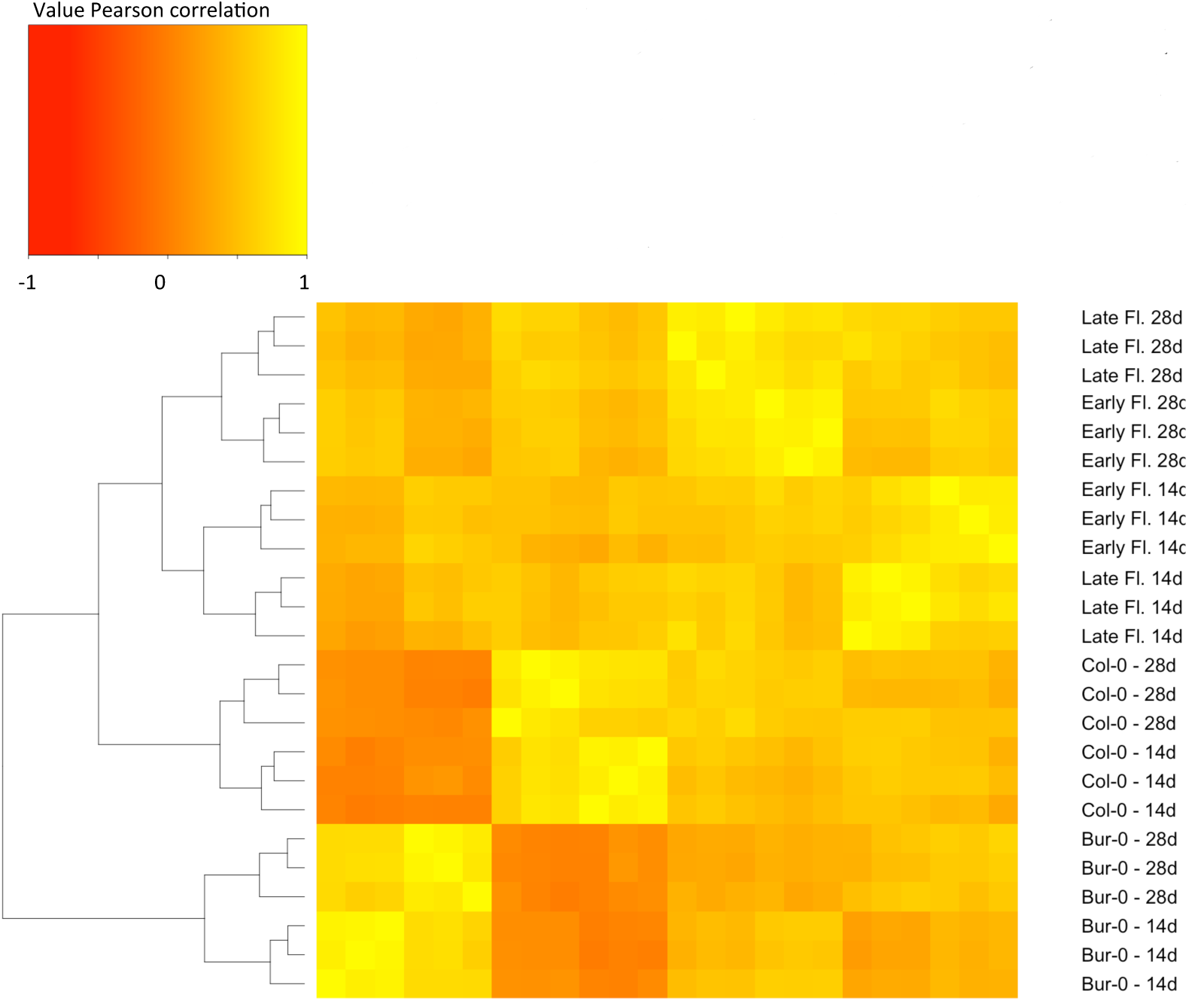
Heatmap of the correlation in gene expression variation for the 1000 genes showing highest expression variation between samples. Clustering of gene expression levels shows that samples partition by genotype (Col-0, Bur-0), sampling time point (14d, 28d), and flowering time (Early Fl.: Early flowering RIL pool, Late Fl.: Late flowering RIL pool).

**Supplementary datasets:** For each expressed genes, Spearman correlation coefficients and their FDR significance presented separately for each dataset (whole Swedish populations, only vernalization-independent Swedish line, species-wide dataset).

## Acknowledgements

This research was supported by the Deutsche Forschung Gesellschaft (DFG) in the realm of SPP1530 grant ME 2742/2--1, and by the European Research Council with Grant 648617 “AdaptoSCOPE”. Raw data has been deposited in NCBI’s Gene Expression Omnibus (Edgar *et al.* 2002) and are accessible through GEO Series accession number GSE97664.

